# Bistable nerve conduction

**DOI:** 10.1101/2022.01.25.477744

**Authors:** Zhaoyang Zhang, Zhilin Qu

## Abstract

The essence of the nerve system is to transmit information from and to different parts of the body in response to environmental changes. Here we show that a bistable conduction behavior can occur in the nerve system, which exhibits stimulus-dependent fast and slow conduction waves. The bistable behavior is caused by a positive feedback loop of the wavefront upstroke speed, mediated by the sodium channel inactivation properties, which is further potentiated by the calcium current. This provides a mechanism for the slow and fast conduction in the same nerve system observed experimentally. We show that the bistable conduction behavior is robust with respect to the experimentally determined activation thresholds of the known sodium and calcium channel families. The theoretical insights provide a generic mechanism for stimulus-dependent fast and slow conduction in the nerve system, which is applicable to conduction in other electrically excitable tissue, such as cardiac muscles.

## Introduction

The major function of the nerve system is to transmit information via electrical excitation and conduction for the body or parts of the body to respond to environmental changes. The response time is determined by the conduction velocity (CV) of the nerve cable. It is well established that CV is determined by the conductance of the sodium (Na^+^) current (I_Na_) and calcium (Ca^2+^) current (I_Ca_), the size of the nerve cable, temperature, and myelination, etc ^1–4^. CV in different nerve fibers or species spans in a wide range, differing in orders of magnitude ^1^. These conduction properties can be well described by the cable equation with the Hodgkin-Huxley (HH) model ^5–10^.

Besides the regular conduction properties, an interesting conduction behavior was observed experimentally, i.e., stimulus-dependent fast and slow conduction waves occur in the same nerve fiber. For example, *Aglantha digitale*, a species of jellyfish, has two modes of swimming—a slow swimming for fishing and a fast swimming for escaping away from predators ^11,12^. In an experimental study ^13^, Mackie and Meech showed that the slow and fast swimming modes were caused by a slow conduction (~0.3 m/s) and a fast conduction (~1.4 m/s) in the same motor giant axon, respectively. The slow conduction is a low-amplitude and long-duration wave mediated by I_Ca_ and the fast conduction is a high-amplitude and short-duration wave mediated by I_Na_ (Fig.S1). The fast wave was induced by an external stimulus and the slow wave was spontaneous and endogenous. This same conduction dynamics was also shown in experiments of conduction in cockroach giant axons by Hochner and Spira ^14^ who demonstrated that two distinct conduction waves (0.1-0.6 m/s and 3-6 m/s) occurred in the same axon treated with ethanol. Differing from *Aglantha digitale*, the slow conduction in the cockroach giant axon was not mediated by I_Ca_ but still by I_Na_. Besides conduction in the giant axons of jellyfish and cockroaches, evidence of stimulus-dependent fast and slow conduction has also been shown in experiments of rat visual cortex ^15,16^, i.e., the CV of the spontaneous waves differ from those of the evoked waves in the same cortex. These experimental observations imply that besides the regular conduction behavior, a new conduction dynamics can occur in the nerve systems, i.e., the same nerve fiber or tissue can exhibit two stable conduction states depending on the initial conditions. As in *Aglantha digitale*, the two conduction behaviors of the same fiber accomplish two distinct biological functions, however, the underlying mechanism remains to be elucidated.

In this study, we investigate the mechanisms of the stimulus-dependent fast and slow conduction using analytical methods and computer simulations in a cable equation with the HH model ^17^. An I_Ca_ formulation is added to the HH model to investigate the role of I_Ca_. We show that stimulus-dependent fast and slow conduction capturing the experimental observations can occur in the cable with the HH model. This is a bistable behavior, namely bistable conduction, emerging during conduction in the cable, caused by a positive feedback loop of the wavefront upstroke speed mediated by the Na^+^ channel inactivation properties. The addition of I_Ca_ can further potentiate bistable conduction. Using simulations of randomly selected parameter sets, we show that the bistable conduction mechanism is robust, i.e., the activation thresholds of I_Na_ and I_Ca_ for bistable conduction detected in a wide range of parameters are well within the experimentally determined activation thresholds of the known Na^+^ and Ca^2+^ channel families. Since the bistable conduction is mediated by the I_Na_ alone in the HH model, it is likely a generic mechanism applicable to conduction not only in the never systems, but also in other electrically excitable media, such as cardiac muscles.

## Results

### Bistable conduction in the cable equation

To observe bistable conduction in the cable equation with the HH model ^17^, we change some of the parameters of the model. To examine the role of I_Ca_, we add an I_Ca_ formulation to the model. The details of the model are presented in Supplemental Information (*SI*). Figs.1 A and B show a fast wave (1.4 m/s) induced by a strong stimulus and a slow wave (0.21 m/s) induced by a weak stimulus, respectively. The action potential of the fast wave (inset in Fig.1A) exhibits a steep upstroke, a high amplitude (≈95 mV, from −65 mV to 30 mV), and a short duration (≈10 ms), while that of the slow wave (inset in Fig.1B) exhibits a shallow upstroke, a low amplitude (≈45 mV, from −65 to −20 mV), and a long duration (≈40 ms). These features recapitulate well the experimental observations in the jellyfish (see Fig.S1) and cockroach experiments.

**Figure 1.**
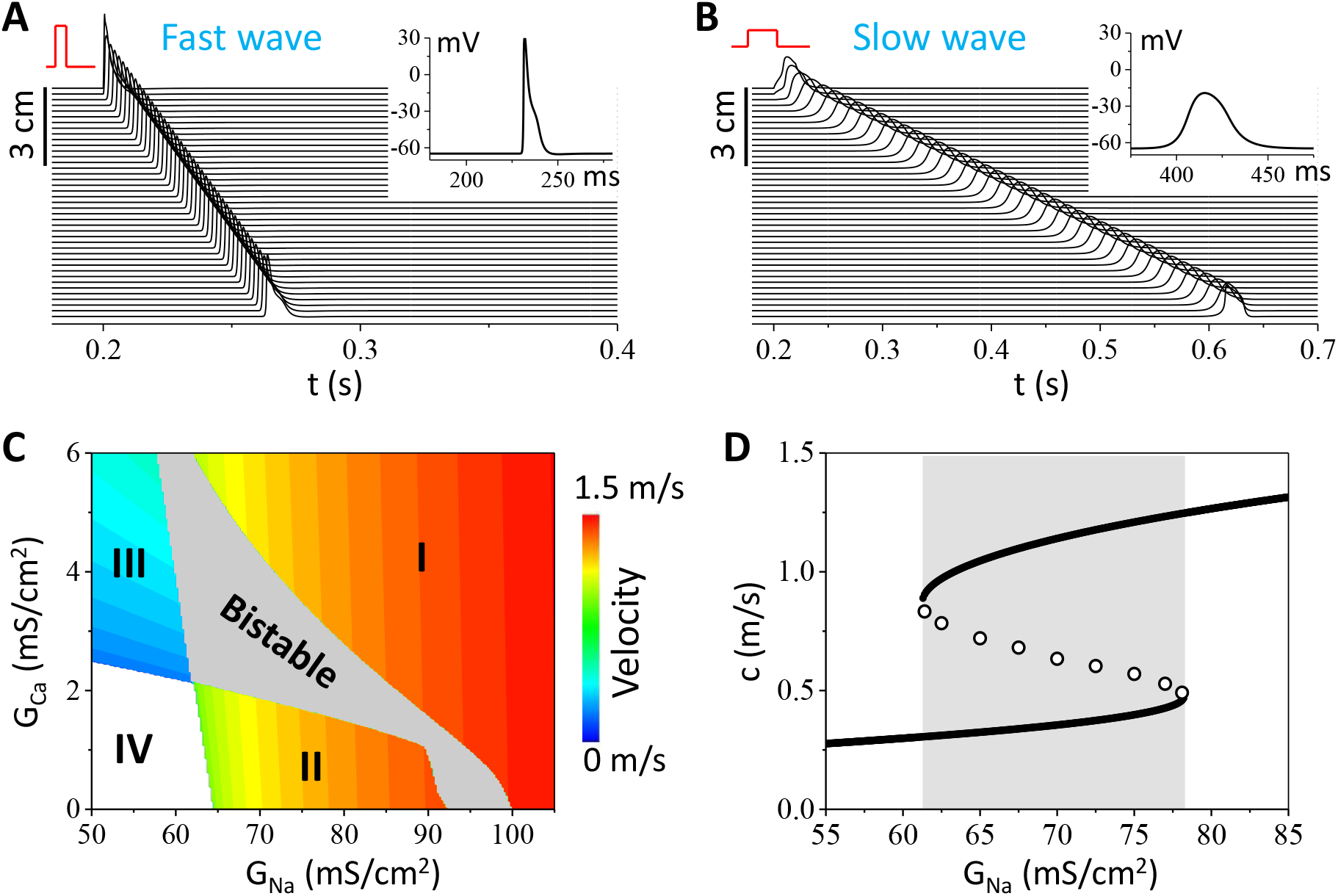
Bistable conduction in the cable equation. **A** and **B**. Stimulus-dependent fast (A) and slow (B) conduction in the same cable. The stimulus in A is 200 μA/cm^2^ with a 0.5 ms duration while the stimulus in B is 5 μA/cm^2^ with a 20 ms duration. The stimulus is applied to the first 0.225 cm of the cable. Insets show action potentials from the middle of the cable for the two cases. G_Na_=95 mS/cm^2^ and G_Ca_=0 mS/cm^2^. **C**. Phase diagram showing conduction behaviors in the G_Na_ and G_Ca_ plane. Regions I and II are monostable fast conduction, region III is monostable slow conduction, region IV is conduction failure, and the gray region is bistable conduction. The phase diagram is obtained using the strong and weak stimulus protocols as in A and B. **D**. CV versus G_Na_ for G_Ca_=3 mS/cm^2^. Solid circles are stable conduction and open circles are unstable conduction (the saddle points). The saddle points are determined by a stimulus very close to the critical stimulus (see Fig.2A for an example). The gray marks the bistable region.

To show how the bistable conduction in the cable is affected by I_Na_ and I_Ca_, we scan the maximum conductance of I_Na_ (G_Na_) and that of I_Ca_ (G_Ca_), and plot the CV in color in the two-parameter plane (Fig.1C). For each parameter set, a strong stimulus and a weak stimulus as in Figs. 1 A and B are applied to elicit conduction waves. The gray region in Fig.1C is where bistable conduction occurs. The white region is where both stimuli fail to elicit conduction (conduction failure). The colored regions (I, II, and III) are where both stimuli give rise to a single stable conduction with CV color coded. Regions I and II exhibit stable fast conduction mediated by I_Na_, and region III exhibits stable slow conduction mediated by I_Ca_. The presence of I_Ca_ promotes bistable conduction, however, I_Ca_ is not required since the bistable conduction occurs in the absence of I_Ca_ (G_Ca_=0). In the absence of I_Ca_ (G_Ca_=0), bistable conduction occurs in an intermediate range of G_Na_, and the fast conduction occurs when G_Na_ is either above or below this range. Therefore, bistable conduction can occur in the HH model mediated by I_Na_ alone without requiring the presence of I_Ca_ or another inward current. In Fig.1D, we plot CV versus G_Na_ for a fixed G_Ca_, showing a typical hysteresis, a hallmark of bistability.

### Mechanism of bistable conduction

Since bistable conduction can occur in the absence of I_Ca_, we investigate how it occurs in in the original HH model, focusing on the role of I_Na_. Fig.2A shows a fast wave induced by a stimulus slightly above the critical strength and a slow wave by a stimulus slightly below the critical strength in the absence of I_Ca_. In the first 40 ms, the action potentials are almost identical for the two cases, which then bifurcate (as indicated by the arrow) into a stable fast wave and a stable slow wave. Since the stimuli are very close to the critical strength, the two waves in the first 40 ms are close to the solution of the unstable conduction. The open circles in Fig. 1D are calculated from the unstable conduction. In the three conduction behaviors, besides the difference in the amplitude and duration of the action potentials, another important difference is the upstroke speed of the wavefront. For the degeneration from the unstable conduction to the stable fast wave, the upstroke speed of the wavefront becomes faster and faster until reaching the steady-state conduction. For the degeneration from the unstable conduction to the stable slow wave, the upstroke speed of the wavefront becomes slower and slower until reaching the steady-state conduction. Because of the different upstroke speeds, the inactivation of I_Na_ is different. In Fig.2B, we plot the steady-state inactivation curve of I_Na_ (h_∞_, thick green) and the trajectories of the three types of conduction in the V-h plane. The trajectory of the steady-state slow wave (thick red) undergoes a path that is very close to the steady-state inactivation curve, while the trajectory of the steady-state fast wave (thick blue) undergoes a path that is far away from the steady-state curve. The thin red and blue trajectories are the transient ones degenerating from the steady-state unstable wave (dashed) to the two stable waves, respectively. In other words, the steady-state unstable trajectory (dashed) is the separatrix of the two stable waves.

**Figure 2.**
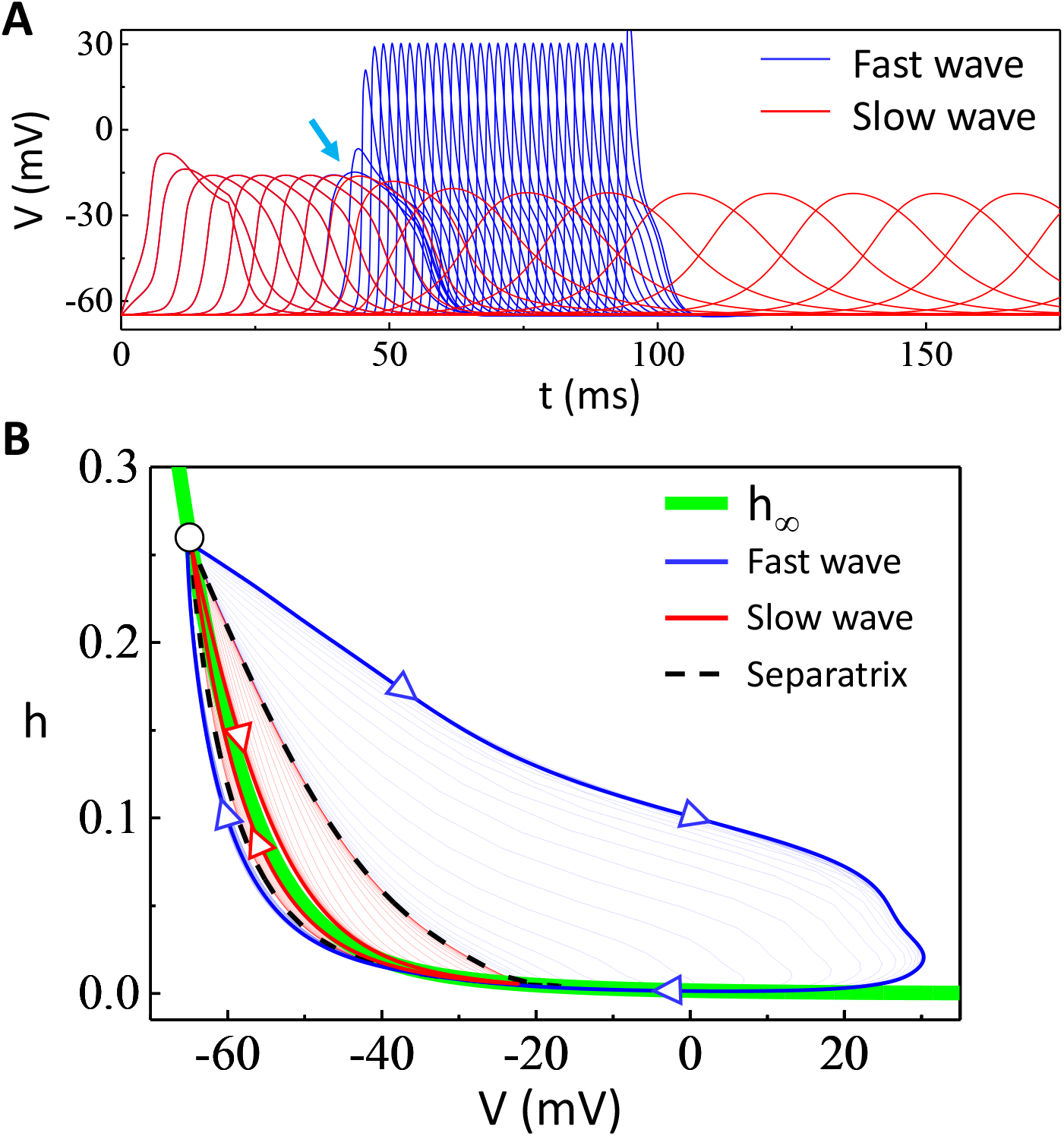
Mechanism of INa-mediated bistable conduction. **A**. Formation of stable fast (blue) and slow (red) conduction from the unstable conduction in the absence of I_Ca_. The fast wave is induced by a stimulus (I_stim_=22.32155910275 μA/cm^2^ with 20 ms in duration) slightly above the critical strength and the slow wave is by a stimulus (I_stim_=22.32155910270 μA/cm^2^ with 20 ms in duration) slightly below the critical strength. Voltage traces are recorded at sites 0.225 cm apart along the cable. The blue and red traces are overlapped until a distance away (marked by the arrow) from the stimulation site. G_Na_=92 mS/cm^2^ and G_Ca_=0. **B**. Na^+^ channel inactivation properties during the fast and slow waves. The thick green line is the steady-state inactivation curve of the Na^+^ channel (h_∞_). The thick red curve with arrows is the trajectory of the steady-state slow wave and the thin red curves are the ones recorded from the transition period from the unstable conduction to the stable slow conduction. The thick blue curve with arrows is the trajectory of the steady-state fast wave and the thin blue curves are the ones recorded from the transition period from the unstable conduction to the stable fast conduction. The black dashed curve is the separatrix of the two types of waves, which is the trajectory taken from the unstable conduction (before the arrow marked in A).

As indicated by the two cases above, a slower upstroke speed causes more I_Na_ inactivation, which in turn results in an even slower upstroke in the next conduction site, or vice versa, forming a positive feedback loop. Therefore, the formation of the stable fast and slow conduction in the cable is a result of the positive feedback in upstroke speed of the wavefront mediated by I_Na_ inactivation, result in two modes of I_Na_ activation/inactivation. This causes bistable conduction to occur in the intermediate range of G_Na_. When G_Na_ is too large, the high I_Na_ mode is always activated and thus only the fast wave occurs. When G_Na_ is too small, the low I_Na_ is not large enough to support the slow wave and thus only the high I_Na_-mediated fast wave is observed. However, I_Ca_ can rescue the slow wave and thus extends the bistable conduction in a wider parameter space (Fig.1C).

To further understand the mechanism of bistable conduction, we obtain an analytical solution using a two-variable model which includes only I_Na_ and I_L_, i.e.,

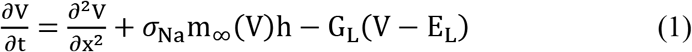

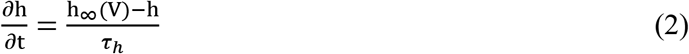

where *V* is voltage and *h* the inactivation gating variable of I_Na_. *σ*_Na_ is a parameter proportional to G_Na_. *m*_∞_(*V*) and *h*_∞_(*V*) are Heaviside functions:

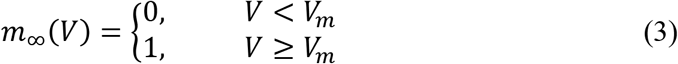

and

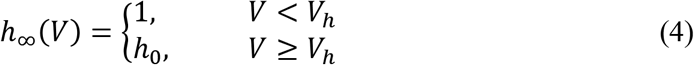

By using the moving coordinate system and continuation conditions, one can obtain that the conduction velocity c satisfies the following equation (see SI method for detailed derivation):

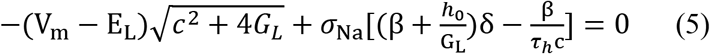

where 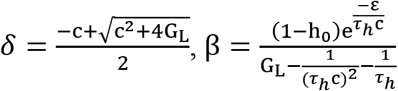, and 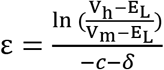. One can rewrite Eq.5 in the form as *σ*_Na_ = *f*(*c*, *τ_h_*, *V_m_*, *V_h_*, *h*_0_, *G_L_*, *E_L_*), i.e., *σ*_Na_ is expressed as a nonlinear function of c. Fig.3A plots c versus *σ*_Na_ calculated using this formulation with other parameters fixed, showing that c is bistable in a certain range of *σ*_Na_. One can also numerically solve Eq.5 to obtain c when the parameters are given. Fig.3B shows the conduction behaviors versus *σ*_Na_ and *τ*_h_, showing that the bistable region decreases as *τ*_h_. We perform additional simulations of the 1D cable with the HH model to investigate the effects of I_Na_ kinetics on the conduction behaviors. Fig.3C shows the conduction behaviors versus *τ*_h_ and *τ*_m_ (activation time constant of I_Na_) and Fig.3D shows the conduction behaviors versus *G*_Na_ and *τ*_h_. Note that the phase diagram in Fig.3D is similar to that in Fig.3B, indicating that theoretical predictions agree well with the numerical simulation results of the HH model.

**Figure 3.**
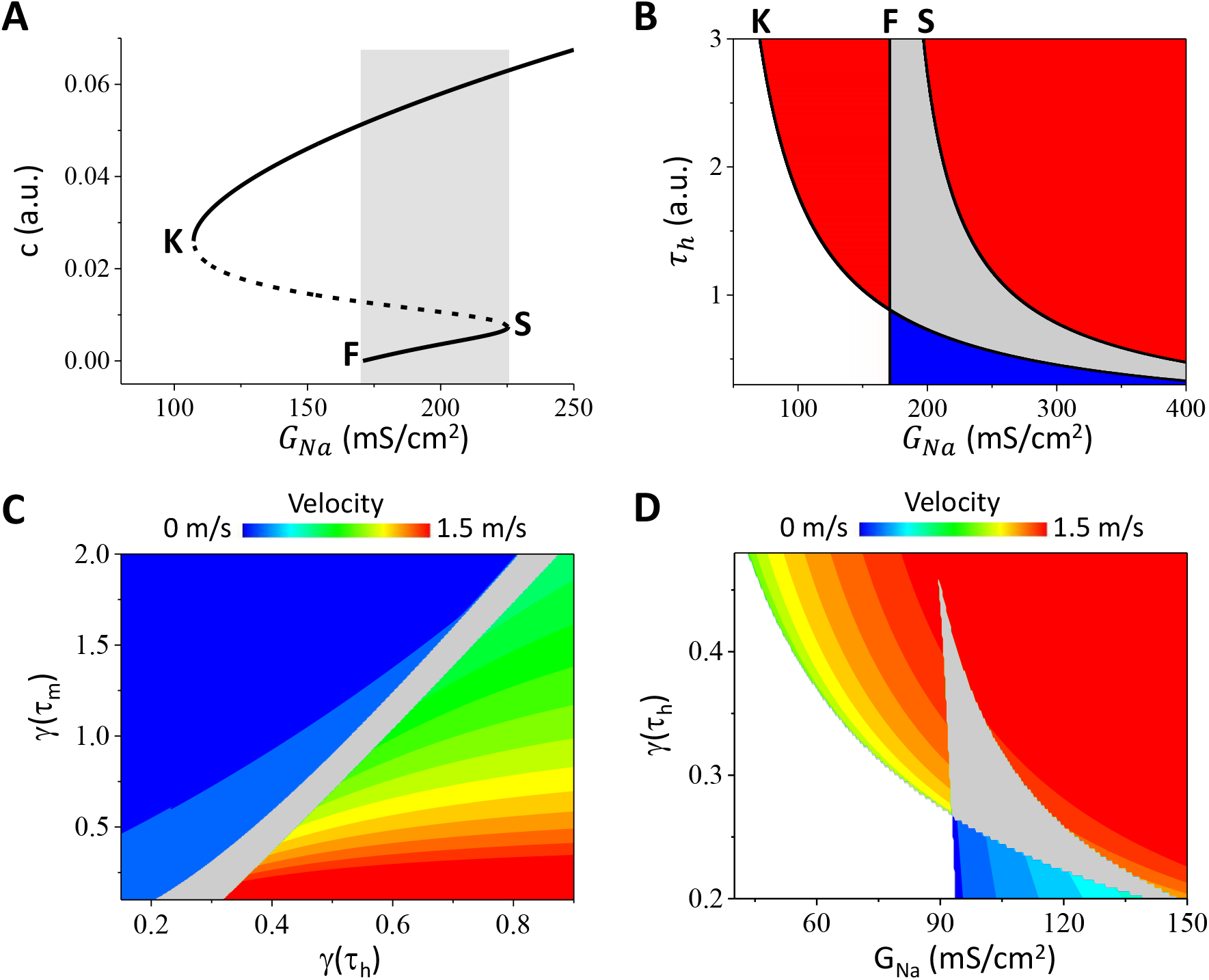
Effects of I_Na_ kinetics on bistable conduction. **A**. Conduction velocity c versus *G_Na_* calculated from the analytical result Eq.5. K and S mark the saddle-node bifurcation points, and F marks the point of conduction failure of the slow conduction. Solid lines are stable conduction and dashed line is unstable conduction. Gray marks the bistable conduction region. *τ_h_* = 1.6, *V_m_* = −19.5 mV, *V_h_* = −55.5 mV, *h*_0_ = 0.075, *G_L_* = 0.3, *E_L_* = −65 mV. We used *σ_Na_* = 2.1G_*Na*_ for Eq.5. **B**. Conduction behaviors versus *G_Na_* and *τ_h_* obtained from the analytical result Eq.5. K, F, and S are the boundaries as marked on A. The gray region is the bistable region. The red regions are monostable fast conduction, the blue region is monostable slow conduction, and the blank region is conduction failure. The parameters are the same as for A. **C**. Conduction behaviors versus *γ*(*τ_h_*) and *γ*(*τ_m_*) from the simulation of the cable equation using the HH model. (γ is the fold change of τ_h_ or τ_m_ from their original values in the HH model, see Eq.S4 in *SI*). The gray region is the bistable conduction region. G_Na_=100 mS/cm^2^ and G_Ca_=0. **D**. Conduction behaviors versus *G_Na_* and *γ*(*τ_h_*) from the simulation of the cable equation using the HH model. Other parameters were the same as for C. The phase diagrams in C and D were obtained and colored the same way as for Fig.1C.

### Robustness of bistable conduction

In the case of *Aglantha digitale* ^13^, based on their observation that the fast wave is mediated by I_Na_ and the slow wave by I_Ca_, the authors hypothesized that to facilitate the fast and slow conduction in the same axon, the Na^+^ channel activation threshold is much higher than the Ca^2+^ channel activation threshold so that during I_Ca_-mediated conduction (the slow wave), the peak voltage remains low enough to avoid activation of I_Na_. On the other hand, in the case of cockroach experiments ^14^, since the slow wave was still mediated by I_Na_, the authors then hypothesized that there might exist another type of I_Na_ with a low activation threshold in the same axon to explain their observations. In other words, theses authors hypothesized that the fast and slow conduction are mediated separately by two inward currents with a certain required difference in activation thresholds (we call it dual-threshold hypothesis). In the original HH model, since there is only one inward current (i.e., I_Na_), it can only exhibit a single-threshold response (Fig.4A). On the other hand, after adding I_Ca_ to the HH model, it can exhibit a dual-threshold response in which two types of action potentials occur depending on the stimulus strength (Fig.4B). Since there are two distinct types of action potentials, one would expect that each type will give rise to a conduction in the cable, resulting in stimulus-dependent fast and slow conduction. This raises a question: which of the two mechanisms is robust or more likely to occur in the real systems?

**Figure 4.**
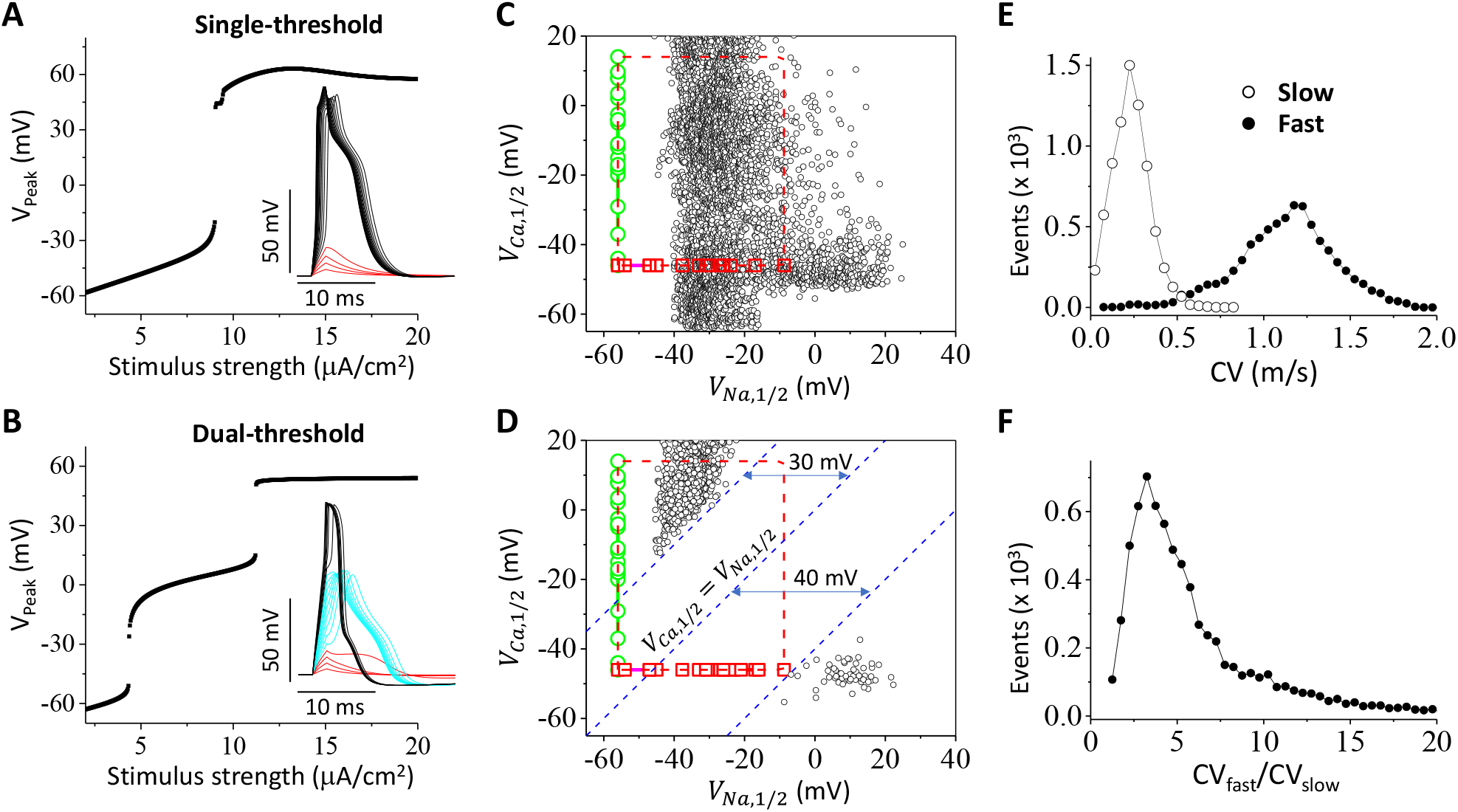
Robustness of bistable conduction. **A**. Peak voltage versus stimulus strength for a case exhibiting a single threshold. Inset shows the voltage traces for different stimulus strengths. Red traces are the subthreshold responses. **B**. Same as A but for a case exhibiting dual thresholds. Inset shows that besides the subthreshold response, there are two distinct types of action potentials. **C**. Parameter sets (black) that give rise to bistable conduction filtered with the single-threshold criterion, plotted in the V_Na,1/2_ and V_Ca,1/2_ plane. Red squares are V_Na,1/2_ taken from Catterall et al ^18^ and green circles are V_Ca,1/2_ taken from Catterall et al ^19^. **D**. Same as C but for the parameter sets filtered with the dual-threshold criterion. Dashed blue lines are reference lines marking the minimum difference of the two activation thresholds. **E**. CV distributions for the slow and fast conduction calculated from the data shown in C. **F**. Distribution of the fast-to-slow CV ratio calculated from the same data in C.

To address this question, we perform the following investigations. We first carry out a large number of simulations of the cable equation with randomly selected parameters in assigned intervals (see details in *SI*). We detect the parameter sets giving rise to both fast and slow conduction. We then use these parameter sets to perform single-cell simulations to filter out the single-threshold and dual-threshold responses by examining the stimulus-response relationship as shown in Figs.4 A and B. The parameter sets (open dots) exhibiting single-threshold responses are plotted in Fig.4C and those exhibiting dual-threshold responses in Fig.4D. The data points are plotted on the plane of the half-activation voltages for Na^+^ channels (V_Na,1/2_) and Ca^2+^ channels (V_Ca,1/2_). See *SI* for definitions and ranges of V_Na,1/2_ and V_Ca,1/2_ in the model. We also plot the experimentally measured V_Na,1/2_ (red squares) and V_Ca,1/2_ (green circles) surveyed from literature by Catterall et al ^18,19^. V_Na,1/2_ for the known 10 members of the Na^+^ channel family ranges from −56 mV to −8.8 mV ^18^, and V_Ca,1/2_ for the 10 members of the Ca^2+^ channel family ranges from −46 mV to 14 mV ^19^. V_Na,1/2_ and V_Ca,1/2_ are plotted in a way so that all possible combinations of the 10 members of the Na^+^ channel family and the 10 members of the Ca^2+^ channel family fall inside the dashed red box. Therefore, we argue that the parameter sets fall inside the box are the ones that can occur in the real system.

The data sets exhibiting single-threshold response fall both inside and outside the dashed box (Fig.4C). We plot the distributions of CV for the fast and slow conduction in Fig.4E, and the ratios of the fast-to-slow CV in Fig.3F. The average slow CV is 0.23 ±0.08 m/s and the average fast CV is 1.10±0.32 m/s. The average of the fast-to-slow CV ratio is 7.0±7.84. The fast-to-slow CV ratios were 5 to 10 in the experimentally measured CVs in the giant axons of *Aglantha digitale* ^13^ and cockroaches ^14^, which are in the same range as obtained in the simulations. Therefore, the bistable conduction mechanism agrees well with experimental data and is robust.

The data sets exhibiting dual-threshold response fall into two groups (Fig.4D). One group is completely outside the box (lower right), which occurs for V_Na,1/2_ is 40 mV higher than V_Ca,1/2_. The other group can still fall into the box (upper left), which occurs for V_Ca,1/2_ is 30 mV higher than V_Na,1/2_. However, some caveats of this group of data are worth noting. The slow wave is mediated by I_Na_ and the fast wave by I_Ca_, which is not what occurs in either the jellyfish ^13^ or the cockroaches ^14^. Although the data sets were filtered through the dual-threshold criterion in single cells, when we check the behaviors in the cable, the Na^+^ channel exhibit similar response as in Fig.2, indicating that the mechanism may still be bistable conduction as in the single-threshold case. For example, when we remove I_Ca_ from the data sets in Fig.4D, we can still observe bistable conduction. To show this, we perform the same simulations using the parameter sets in Fig.4D without I_Ca_ (See Fig.S2A), the parameter sets outside the box disappear but about 50% of the parameter sets inside the box retain. Another evidence to support this is that if we slow the Na^+^ channel inactivation, this group of parameter sets will disappear. We perform the same simulations as in Fig.4 except that the inactivation is slowed by increasing *γ*(*τ_h_*) = 0.35 to *γ*(*τ_h_*) = 1 (see Fig.S2B). The upper group disappears, but the lower group retains and is still outside the dashed box. The results shown in Fig.4 and Fig.S2 indicate that the dual-threshold mechanism is difficult to be satisfied in the real systems.

In the model, the resting potential is set at −65 mV. If we lower the resting potential, e.g., to −75 mV, we observe almost the same results (Fig.S3) as those shown in Figs. 4 and S2 except that the minimum I_Na_ or I_Ca_ activation threshold for bistable conduction has a roughly 10 mV shift due to the lowering of the resting potential.

## Discussion

Nerve conduction can be well described by the cable equation with the HH model ^5–9^. It is well known that the cable equation with FitzHug-Nagumo model or the HH model exhibit monostable conduction ^10^. In this study, we show that the cable equation with the HH model can exhibit bistable conduction in which a fast and a slow stable conduction occur in the same cable depending on the stimulus strength. The bistable behavior is a result of the positive feedback of the wavefront upstroke speed mediated by the Na^+^ channel inactivation properties. In other words, this positive feedback cause two stable modes of I_Na_ activation, which occurs when the Na^+^ channel inactivation is relatively fast (see also Fig.3). Unlike the fast conduction which occurs as long as the G_Na_ is greater than a critical value, bistable conduction can only occur in an intermediate range of G_Na_. This is because when G_Na_ is too large, the high I_Na_ mode is always activated and thus only the fast wave can occur. When G_Na_ is too small, the low I_Na_ cannot support a stable slow conduction, and thus only the fast conduction can occur. However, the failed slow conduction can be rescued by the addition of I_Ca_, which can substantially extend the bistable conduction regime (see Fig.1C). We use an analytical treatment of simplified cable model to demonstrate that bistable conduction can be mediated a single inward current, namely I_Na_. Using simulations with randomly drawing parameter sets, we also show that the bistable conduction is robust, which can occur well within the experimentally determined ranges of the activation thresholds of the known Na^+^ and Ca^2+^ channel families.

Our computer simulation results agree well with the experimental observations of electrical conduction in the giant axons of jellyfish and cockroaches. For example, the action potential profiles in the fast and slow waves and the ratios of the fast-to-slow CV in the simulations agree with those shown in experiments in both jellyfish and cockroaches. More importantly, our theoretical study unifies the seemly different experimental observations in jellyfish and cockroaches to the same general mechanism. In the jellyfish experiments, Mackie and Meech ^13^ showed that the slow wave was blocked by Ca^2+^ channel blockers, and thus concluded that the slow wave was mediated by I_Ca_. This led them to hypothesize that the activation threshold of I_Na_ has to be much higher than that of I_Ca_ to allow the two waves to occur. As shown in our simulations (Fig.1C), although I_Ca_ is not required for bistable conduction, it potentiates bistable conduction by rescuing the slow conduction. Therefore, blocking it will suppress the slow conduction, agreeing with the experimental observation. On the other hand, in the cockroach experiments, Hochner and Spira ^14^ found that the slow wave was blocked not by Ca^2+^ channel blockers but by Na^+^ channel blockers, differing from the jellyfish experiments. They then hypothesized that a low-threshold I_Na_ must be responsible for the slow wave. Our simulations showed that I_Ca_ or another low-threshold I_Na_ is not required since a single I_Na_ could produce both the slow and fast conduction via the bistable dynamics. In other words, the fast and slow conduction observed in cockroaches may originated from the same I_Na_. A moderate I_Na_ reduction may block the slow wave but not the fast wave while a strong I_Na_ reduction can block both waves (see Fig.1C). Therefore, the experimental observations in both jellyfish and cockroaches can be explained by the same mechanism in different parameter regimes.

Besides the giant axons in jellyfish and cockroaches, bistable conduction may also occur in other nerve systems. For example, it was shown that the spontaneous waves and the evoked waves exhibit distinct CVs in the same rat visual cortex ^15,16^, which could be a result of bistable conduction. In addition, bimodal CV distributions were widely observed in sensory and motor nerves ^20–26^, which were traditionally attributed to the size difference of the fibers. However, it was also shown that the action potential profiles in the fast conduction are different from those in the slow conduction, i.e., the amplitude is much lower and the duration is much longer in the slow conduction ^20^, similar to those observed in the jellyfish and cockroaches. The large difference in action potential properties cannot be simply attributed to the size difference of the nerve fibers since although the size of the nerve fibers can largely affect CV, it may only have a small effect on the action potential properties. Bistable conduction can be a candidate mechanism for bimodal CV distributions, which needs to be verified in future experimental studies.

Finally, as shown in our simulations (Fig. 4), the dual-threshold mechanism is theoretically plausible, however, it requires a very large minimum difference in the activation thresholds of I_Na_ and I_Ca_, which may not be satisfied easily in the real systems. On the other hand, the bistable conduction mechanism can be easily achieved without requiring any gap or correlation between the activation thresholds of the two types of ionic currents. Moreover, the bistable conduction mechanism may also have an evolutionary advantage over the dual-threshold mechanism since a single rather than two inward currents can provide two survival functions as in the jellyfish. Although it is unclear what are the functional roles of the fast and slow conduction in other species or diseases, as it is well known that bistability is a ubiquitous phenomenon in biology and responsible for many biological functions ^27–30^, we believe that bistable conduction may also play important roles in nerve functions under health and diseased conditions. As shown in our study, the bistable conduction occurs in the cable equation with the HH model, we believe that it is a generic mechanism that is applicable to not only the nerve systems but also other electrically excitable tissue, such as cardiac muscles.

## Methods

The HH model ^17^ with modifications is used to simulate action potential conduction in a cable with the following partial differential equation for voltage (*V*):

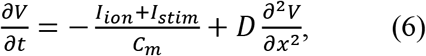

where *C*_m_=1 μF/cm^2^ is the membrane capacitance, and D=0.0045 cm^2^/ms is the diffusion constant. *I_stim_* is the stimulus current density and *I_ion_* is the total ionic current density consisting of different types of ionic currents, i.e.,

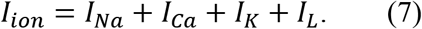

I_Na_ is the Na^+^ current, I_Ca_ is the Ca^2+^ current, I_K_ is the K^+^ current, and I_L_ is the leak current. The formulations of I_Na_, I_K_, and I_L_ are the same as in the HH model. I_Ca_ is formulated based on Medlock et al ^31^ with the addition of an inactivation gate. Eq.1 is numerically solved using a forward Euler method with Δ*x* = 0.045 *cm* and Δ*t* = 0.005 *ms*. The gating variables are integrated using the method by Rush and Larsen ^32^. No-flux boundary condition is used. The mathematical details of the model and numerical algorithms are presented in *SI*.

## Supporting information

Supplemental Methods and Results

## Author contributions

ZQ conceived the project, supervised the research, provided funding, and wrote the manuscript; ZZ performed the simulations and mathematical analyses; ZQ and ZZ analyzed the results and edited the manuscript.

