## Supplemental Methods and Results for "Bistable nerve conduction"

### Supplemental Information

#### A. Supplemental Methods

##### 1. Mathematical model

We used the Hodgkin-Huxley (HH) model (*1*) with modifications to simulate the action potential conduction in a cable with the following partial differential equation for voltage ($V$):

$$\frac{\partial V}{\partial t} = -\frac{I_{ion}+I_{stim}}{C_m} + D\frac{\partial^2 V}{\partial x^2} \qquad \text{(S1)}$$

where $C_m$=1 μF/cm$^2$ is the membrane capacitance, and D=0.0045 cm$^2$/ms is the diffusion constant. $I_{stim}$ is the stimulus current density and $I_{ion}$ is the total ionic current density consisting of different types of ionic currents, i.e.,

$$I_{ion} = I_{Na} + I_{Ca} + I_K + I_L \qquad \text{(S2)}$$

In Eq.S2, $I_{Na}$ is the $Na^+$ current density described by $I_{Na} = G_{Na}m^3h(V - E_{Na})$. $I_K$ is the $K^+$ current density described by $I_K = G_K n^4(V - E_K)$. $I_L$ is the leak current density described by $I_L = G_L(V - E_L)$. $I_{Ca}$ is the $Ca^{2+}$ current density described by $I_{Ca} = G_{Ca}\mathrm{d}^2 f(V - E_{Ca})$. This formulation was adopted from Medlock et al (*2*) with the addition of an inactivation gate *f*. *m*, *h*, *n*, *d* and *f* are gating variables which are described by the following type of differential equations:

$$\frac{dy}{dt} = (y_\infty - y)/\tau_y \qquad \text{(S3)}$$

In Eq.3, $y_\infty = \frac{\alpha_y}{\alpha_y+\beta_y}$ and $\tau_y = \frac{1}{\alpha_y+\beta_y}$ in which $\alpha$ and $\beta$ are rate constants and functions of V.

To observe bistable conduction, we made modifications to the original HH kinetics as detailed in the Table below:

| | **Original HH kinetics** | **Modified HH kinetics** |
|---|---|---|
| $\alpha_m$ | $0.1\frac{25-\mathrm{V}}{\exp(\frac{25-\mathrm{V}}{10})-1}$ | $0.1\frac{-35-\mathrm{V}}{\exp(\frac{-35-\mathrm{V}}{10})-1}$ |
| $\beta_m$ | $4\exp(\frac{-\mathrm{V}}{18})$ | $4\exp(\frac{-60-\mathrm{V}}{18})$ |
| $\alpha_h$ | $0.07\exp(\frac{-\mathrm{V}}{20})$ | $0.07\exp(\frac{-75-\mathrm{V}}{20})$ |
| $\beta_h$ | $\frac{1}{\exp\left(\frac{30-\mathrm{V}}{10}\right)+1}$ | $\frac{1}{\exp\left(\frac{-45-\mathrm{V}}{10}\right)+1}$ |
| $\alpha_n$ | $0.01\frac{10-\mathrm{V}}{\exp(\frac{10-\mathrm{V}}{10})-1}$ | $0.01\frac{-15-\mathrm{V}}{\exp(\frac{-15-\mathrm{V}}{10})-1}$ |
| $\beta_n$ | $0.125\exp(\frac{-\mathrm{V}}{80})$ | $0.125\exp(\frac{-25-\mathrm{V}}{80})$ |
| $\mathbf{E_{Na}}$ | 120 mV | 55 mV |
| $\mathbf{E_K}$ | -12 mV | -77 mV |
| $\mathbf{E_L}$ | 10.6 mV | -65 mV |

The changes were shifts of the voltage, which was to give rise to: 1) a resting potential of -65 mV (0 mV in the original HH model); and 2) bistable conduction. Besides the voltage shifts, we also altered the magnitudes of the time constants $\tau_y$ by multiplying a pre-factor $\gamma(\tau_y)$, i.e.,

$$\tau_y(V) \rightarrow \gamma(\tau_y) \times \tau_y(V) \qquad \text{(S4)}$$

For $d_\infty$ and $f_\infty$, we used the following formulations: $d_\infty = \frac{1}{1+\exp\left(-\frac{V+14}{k_d}\right)}$ and $f_\infty = \frac{1}{1+\exp\left(\frac{V+44}{k_f}\right)}$. $\tau_d$ and $\tau_f$ were set as constants independent of V.

The default parameters were set as: $G_K$=36 mS/cm$^2$, $G_L$=0.3 mS/cm$^2$, $\gamma(\tau_m) = 0.2$; $\gamma(\tau_h) = 0.35$, $\gamma(\tau_n) = 3$; $\tau_d = 3$ ms, $\tau_f = 20$ ms, $k_d = 5.8$, and $k_f = 6$. The values of $G_{Na}$ and $G_{Ca}$ were stated in the figure legends. The parameter intervals for the simulations of random parameter selection were described below.

#### 2. Assigned intervals for random parameter drawing

To evaluate the robustness of the two mechanisms, we carried out a large number of simulations with randomly drawn parameter sets. In these simulations, besides drawing the maximum conductance and time constants randomly, we also randomly shifted the kinetics of the ionic currents to investigate the effects of their activation thresholds on conduction. To shift the $I_{Na}$ kinetics, we replaced V for $\alpha_m$, $\beta_m$, $\alpha_h$, and $\beta_h$ by ($V+V_{Na,shift}$). In the control, the half-activation potential for $I_{Na}$ is $V_{Na,1/2} \approx -35\ mV$ (at which $m_\infty$=0.5). After the shift, the half-activation potential becomes,

$$V_{Na,1/2} = -35 - V_{Na,shift} \qquad \text{(S5)}$$

Similarly, we shifted the $I_{Ca}$ kinetics by replacing V for $d_\infty$ and $f_\infty$ with ($V+V_{Ca,shift}$). The half-activation voltage for $I_{Ca}$ at control is $V_{Ca,1/2} = -14\ mV$ (at which $d_\infty$=0.5), and after the shift, it becomes,

$$V_{Ca,1/2} = -14 - V_{Ca,shift} \qquad \text{(S6)}$$

We also shifted the $I_K$ kinetics by replacing V for $\alpha_n$ and $\beta_n$ with ($V+V_{K,shift}$).

The randomly drawn parameter sets used for the simulations for Fig.3, Fig.4, and Fig.S2 were uniformly drawn from the following pre-assigned intervals: $G_{Na}\in$[50,200], $G_K\in$[20,60], $G_{Ca}\in$[0,10], $V_{Na,shift}\in$[-75,25], $V_{Ca,shift}\in$[-50,50], $V_{K,shift}\in$[-40,60], $\gamma(\tau_m)\in$[0.1,0.5], $\gamma(\tau_d)\in$[1,8], $\gamma(\tau_n)\in$[0.5,10], $k_f\in$[8,2]. The fixed parameters were the same as for control. Therefore, based on Eqs.S5 and S6 and the assigned intervals for $V_{Na,shift}$ and $V_{Ca,shift}$, the range for $V_{Na,1/2}$ is [-60, 40] mV and that for $V_{Ca,1/2}$ is [-64, 36] mV.

#### 3. Numerical simulation methods

Forward Euler method was used for numerical simulation of Eq.S1 with the following discretization:

$$V_i(t+\Delta t) = V_i(t) + \{-\frac{I_{ion}+I_{stim}}{C_m} + D\frac{[V_{i+1}(t)+V_{i-1}(t)-2V_i(t)]}{\Delta x^2}\}\Delta t \qquad \text{(S7)}$$

with $\Delta x = 0.045\ cm$ and $\Delta t = 0.005\ ms$. No-flux boundary condition was used. The gating variables (Eq.S3) were integrated using the method by Rush and Larsen (*3*), i.e.,

$$y(t+\Delta t) = y_\infty - [y_\infty - y(t)]e^{-\Delta t/\tau_y} \qquad \text{(S8)}$$

#### 4. Theoretical model and analysis of bistable conduction

To understand the mechanism of bistable conduction mediated by $Na^+$ channel, we investigated the following simple cable equation:

$$\frac{\partial V}{\partial t} = \frac{\partial^2 V}{\partial x^2} + \sigma_{Na} m_\infty(V) h - G_L(V - E_L) \qquad \text{(S9)}$$

$$\frac{\partial h}{\partial t} = \frac{h_\infty(V) - h}{\tau_h} \qquad \text{(S10)}$$

For simplicity and analytical treatment, we set the diffusion constant D=1, and set $\sigma_{\mathrm{Na}} = \mathrm{G_{Na}}(\mathrm{E_{Na}} - \mathrm{V_C})$ to be single parameter. We also assume $\tau_h$ is a constant independent of V. $m_\infty(V)$ and $h_\infty(V)$ are Heaviside functions of V as follows:

$$m_\infty(V) = \begin{cases} 0, & V < V_m \\ 1, & V \geq V_m \end{cases} \qquad \text{(S11)}$$

and

$$h_\infty(V) = \begin{cases} 1, & V < V_h \\ h_0, & V \geq V_h \end{cases} \qquad \text{(S12)}$$

We seek a travelling wave solution $\mathrm{V(x,t)}$ with speed c in an infinite spatial domain: $x \in (-\infty, +\infty)$, which becomes a stationary front in the moving coordinate system (see Fig.S4) $\varepsilon = x - ct$, i.e.,

$$V(x,t) = U(x - ct) = U(\varepsilon) \qquad \text{(S13)}$$

By substituting Eq.S13 into Eq.S9 and Eq.S10, one obtains the follow set of differential equations:

$$0 = U_{\varepsilon\varepsilon} + cU_\varepsilon + \sigma_{Na} m_\infty h - G_L(U - E_L) \qquad \text{(S14)}$$

$$h_\varepsilon = \frac{h_\infty(U) - h}{-c\tau_h} \qquad \text{(S15)}$$

in which $U_\varepsilon = \frac{dU}{d\varepsilon}$ and $U_{\varepsilon\varepsilon} = \frac{dU_\varepsilon}{d\varepsilon}$. The solution U(ε) satisfies the following boundary conditions: 1) $U_\varepsilon = U_{\varepsilon\varepsilon} = 0$ at $\varepsilon \to \pm\infty$; and 2) We assume that $m_\infty(V)$ changes from 1 to 0 at $\varepsilon = 0$, then $U(0) = V_m$ and $U_\varepsilon|_{\varepsilon=0^-} = U_\varepsilon|_{\varepsilon=0^+}$. Therefore, at the boundaries, U(ε) satisfies:

$$U(\varepsilon) = \begin{cases} U_0, & \varepsilon \to -\infty \\ V_m, & \varepsilon = 0 \\ E_L, & \varepsilon \to +\infty \end{cases} \qquad \text{(S16)}$$

where $U_0 = \frac{\sigma_{Na} h_0 + G_L E_L}{G_L}$. Now the problem is to solve the linear differential equations in the subdomains of $\varepsilon < 0$ and $\varepsilon > 0$.

(i) $\varepsilon > 0$

Since $m_\infty = 0$ when $\varepsilon > 0$, then Eq.S14 becomes:

$$0 = U_{\varepsilon\varepsilon} + cU_\varepsilon - G_L(U - E_L) \qquad \text{(S17)}$$

which has the following form of solution:

$$U(\varepsilon) = Ae^{\lambda_- \varepsilon} + E_L \qquad \text{(S18)}$$

where $\lambda_- = \frac{-c-\sqrt{c^2+4G_L}}{2}$ is a solution of the characteristic equation $\lambda^2 + c\lambda - G_L(U - E_L) = 0$. Since $\lambda_- < 0$, $U(\varepsilon) = E_L$ at $\varepsilon \to +\infty$ is satisfied. Since $U(\varepsilon) = V_m$ at $\varepsilon = 0$, one obtains $A = V_m - E_L$. The solution of Eq.S15 in the whole spatial domain is:

$$h(\varepsilon) = \begin{cases} h_0 + (1 - h_0)e^{\frac{\varepsilon-\varepsilon_0}{c\tau_h}}, & \varepsilon < \varepsilon_0 \\ 1, & \varepsilon \geq \varepsilon_0 \end{cases} \qquad \text{(S19)}$$

in which $\varepsilon_0$ is determined by $(\mathrm{V_m} - \mathrm{E_L})\mathrm{e}^{\lambda_-\varepsilon_0} + \mathrm{E_L} = V_h$, which leads to

$$\varepsilon_0 = \frac{ln\,(\frac{V_h-E_L}{V_m-E_L})}{\lambda_-} \qquad \text{(S20)}$$

(ii) $\varepsilon < 0$

Since $m_\infty = 1$ when $\varepsilon < 0$, using Eq.S19 for h, then Eq.S14 becomes

$$0 = U_{\varepsilon\varepsilon} + cU_\varepsilon + \sigma_{Na}[h_0 + (1 - h_0)e^{\frac{\varepsilon-\varepsilon_0}{c\tau_h}}] - G_L(U - E_L) \qquad \text{(S21)}$$

Eq.S21 exhibits the following form of solution:

$$U(\varepsilon) = U_0 + Be^{\lambda_+\varepsilon} + \beta e^{\frac{\varepsilon}{c\tau_h}} \qquad \text{(S22)}$$

where $\lambda_+ = \frac{-c+\sqrt{c^2+4G_L}}{2}$ is a solution of the characteristic equation $\lambda^2 + c\lambda - G_L(U - E_L) = 0$. Since $\lambda_+ > 0$, $U(\varepsilon) = U_0$ at $\varepsilon \to -\infty$ is satisfied. Since $U(\varepsilon) = V_m$ at $\varepsilon = 0$, one obtains $B = V_m - \beta - U_0$. Inserting Eq.S22 into Eq.S21, one obtains the following equation for β:

$$\frac{\beta}{(c\tau_h)^2} + \frac{c\beta}{c\tau_h} + \sigma_{Na}(1 - h_0)e^{\frac{-\varepsilon_0}{c\tau_h}} - \beta G_L = 0 \qquad \text{(S23)}$$

which gives rise to $\beta = \frac{\sigma_{Na}(1-h_0)e^{\frac{-\varepsilon_0}{c\tau_h}}}{G_L - \frac{1}{(c\tau_h)^2} - \frac{1}{\tau_h}}$.

Finally, applying the constrain that U is smooth at $\varepsilon = 0$, i.e., $U_\varepsilon|_{\varepsilon=0^-} = U_\varepsilon|_{\varepsilon=0^+}$, one has from Eq.S18 and Eq.S22 the following relation:

$$\mathrm{A}\lambda_- = \mathrm{B}\lambda_+ + \frac{\beta}{c\tau_h} \qquad \text{(S24)}$$

which leads to

$$-(\mathrm{V_m} - \mathrm{E_L})\sqrt{c^2 + 4G_L} + \sigma_{\mathrm{Na}}[(\beta + \frac{h_0}{\mathrm{G_L}})\lambda_+ - \frac{\beta}{\tau_h c}] = 0 \qquad \text{(S25)}$$

Therefore, once the parameters are given, one can solve Eq.S25 to obtain the conduction speed c as a function of anyone of the parameters, which is discussed in the main text. Fig.S4 plots the two stable and the unstable traveling wave solutions of Eq.S18 and Eq.S22 with the c values calculated from Eq.S25.

#### B. Supplemental Figures

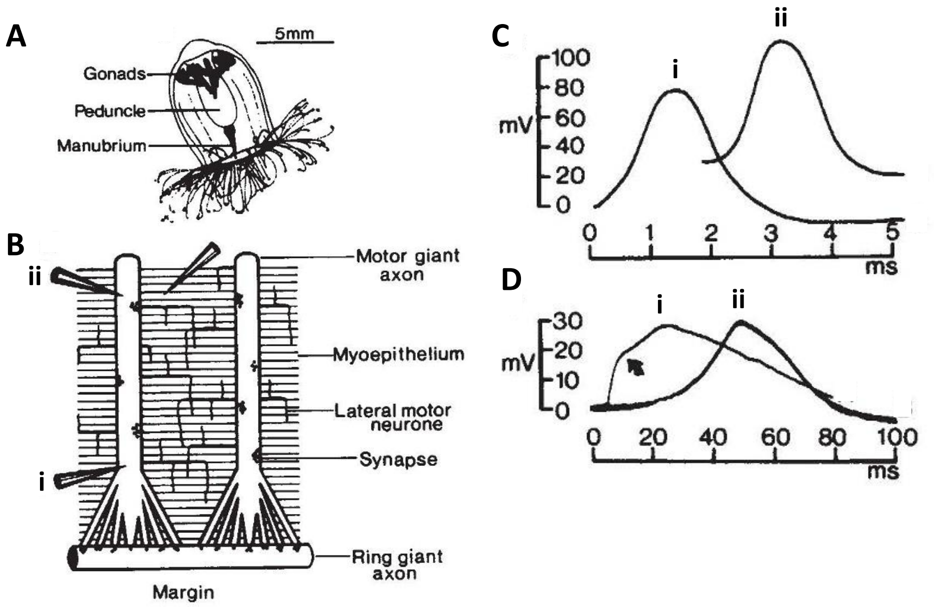


**Figure S1. Action potential conduction in giant axon of *Aglantha digitale*. A**. Schematic drawing of *Aglantha digitale*. **B**. Recording sites (i and ii) in the giant axon. **C**. Recorded action potentials during an escape swim, which are $I_{Na}$-mediated high amplitude and short duration excitations. **D**. Recorded action potentials during slow swim, which are $I_{Ca}$-mediated low amplitude and long duration excitations. This figure is modified from Mackie and Meech (*4*), see the original paper for a detailed description.

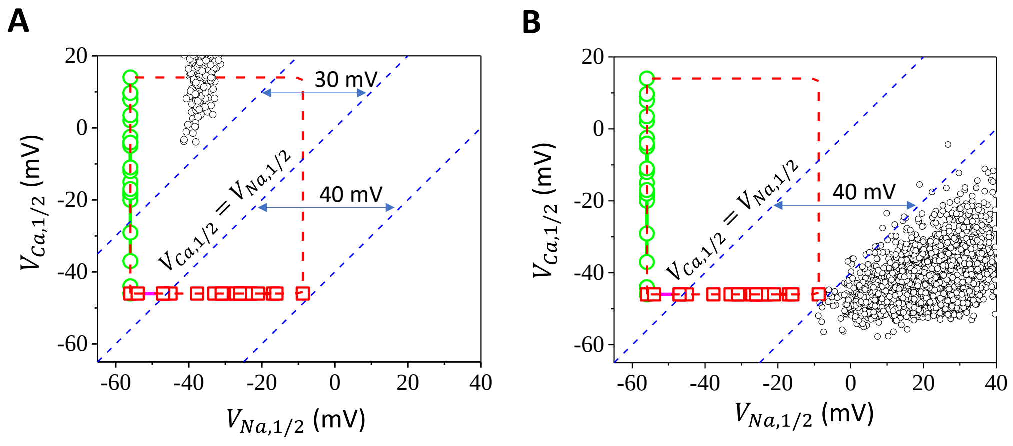


**Figure S2**. **Robustness of bistable conduction. A**. Parameter sets (black) exhibiting stimulus-dependent fast and slow conduction after $I_{Ca}$ was removed from the model using the parameter sets in Fig.4D. The data points in the lower-right group are gone but there are data points in the upper-left group. This indicates that the upper-left group in Fig.4D may still undergo the bistable conduction even though the parameter sets were filtered using the dual-threshold criterion. **B**. Parameter sets (black) exhibiting stimulus-dependent fast and slow conduction after changing $\gamma(\tau_h)$=0.35 to $\gamma(\tau_h)$=1. Unlike Figs. 4 C or D, the data points in this panel were not filtered with the threshold criterion.

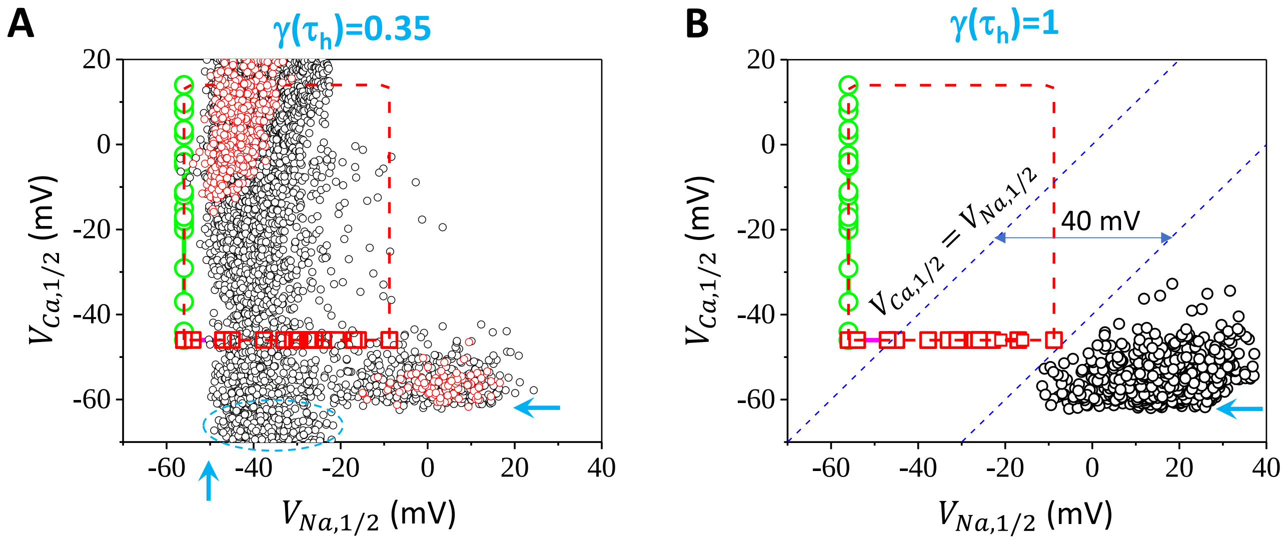


**Figure S3**. **Effects of lowering the resting potential**. Data sets exhibiting fast and slow conduction after changing the resting potential from -65 mV to -75 mV by changing $E_L$=-65 mV to $E_L$=-75 mV. **A**. $\gamma(\tau_h)$=0.35. Red dots are the parameter sets filtered with the dual-threshold criterion. Black dots are the parameter sets filtered with the single-threshold criterion. **B**. $\gamma(\tau_h)$=1. Simulation protocols and parameters are the same as for Fig.3 in the main text except the mentioned changes. The lowering of the resting potential caused two changes: down-shift of the $V_{Ca,1/2}$ boundaries (marked by horizontal arrows, ~10 mV lower comparing to Fig.3 and Fig.4) and left-shift of the $V_{Na,1/2}$ boundary in A (marked by the vertical arrow, ~10 mV left-shifted comparing to Fig.3C). In other words, shifting the resting from -65 mV to -75 mV results in a ~10 mV shift in these boundaries. Beyond these boundaries, spontaneous oscillations occur and thus no conduction in the cable due to the low activation thresholds. The data points in the region marked by the dashed circle are the bistable conduction caused by $I_{Na}$ alone in the absence of or under weak $I_{Ca}$ so that the lower boundary becomes lower than the oscillation boundary caused by $I_{Ca}$.

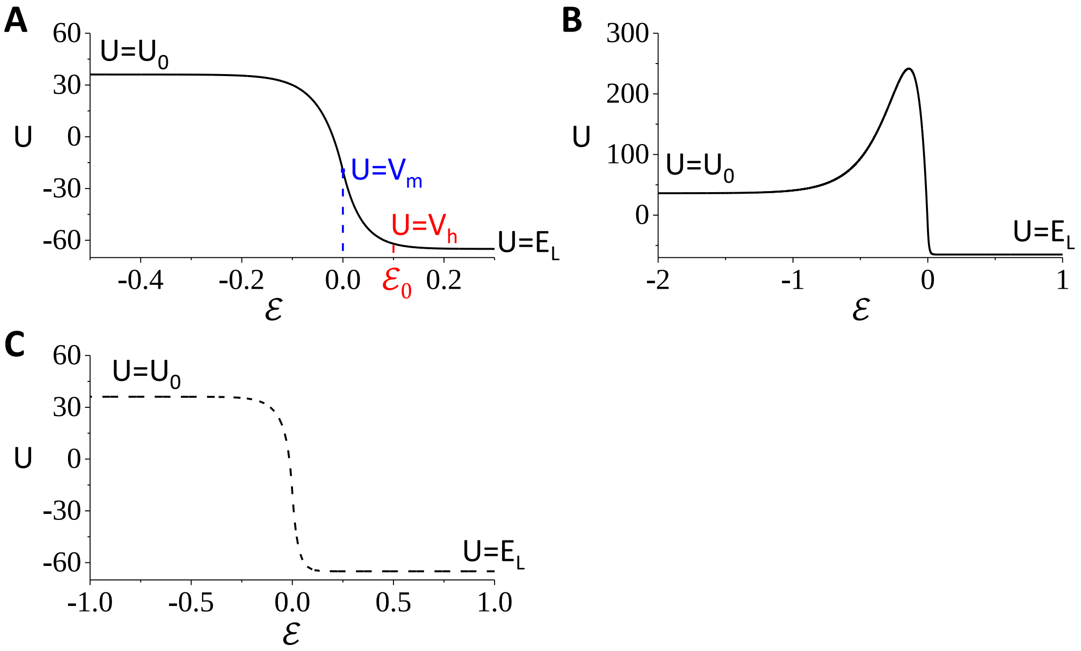


**Figure S4**. **Stable and unstable wavefronts in the moving coordinate from the simplified model**. **A**. The stable slow wave front of a bistable conduction. U=$V_m$ when ε=0, which gives rise to $m_\infty$=1 when ε<0 and $m_\infty$=0 when ε>0. U=$V_h$ when ε=$\varepsilon_0$, which gives rise to $h_\infty$=$h_0$ when ε<$\varepsilon_0$ and $h_\infty$=1 when ε>$\varepsilon_0$. **B**. The stable fast wave front of the same bistable conduction as in A. **C**. The unstable wave front of the same bistable conduction as in A and B. The traveling wave solutions in A-C were plots of Eq.S18 and Eq.S22 using c values calculated in Eq.S25 with the following parameter set: $\tau_h = 1.6$, $V_m = -19.5$ mV, $V_h = -55.5$ mV, $h_0 = 0.075$, $G_L = 0.3$, $E_L = -65$ mV. $G_{Na} = 190$ mS/cm$^2$. $\sigma_{Na} = 2.1G_{Na}$.
